# Random barcode transposon-site sequencing in *Mycobacterium tuberculosis* reveals the functions of uncharacterized genes

**DOI:** 10.1101/2025.11.13.688195

**Authors:** Kayla M. Dinshaw, Katie A. Lien, Matthew Knight, Sorel V. Yigma Ouonkap, Hualan Liu, David F. Savage, Hans K. Carlson, Adam M. Deutschbauer, Sarah A. Stanley

**Affiliations:** Department of Molecular and Cell Biology, University of California, Berkeley, Berkeley, California, USA; Department of Plant and Microbial Biology, University of California, Berkeley, Berkeley, California, USA; Joint Genome Institute, Lawrence Berkeley National Laboratory, Berkeley, California, USA; Howard Hughes Medical Institute, University of California, Berkeley, Berkeley, CA; Environmental Genomics and Systems Biology Division, Lawrence Berkeley National Laboratory, Berkeley, California, USA

## Abstract

*Mycobacterium tuberculosis* (Mtb) is a human bacterial pathogen that establishes chronic infection in the lung. Although the genome of Mtb was sequenced nearly 25 years ago, the genetic basis of Mtb’s success as a human pathogen remains to be fully elucidated. Large-scale genetic approaches to understanding gene function are hindered by the limited throughput of traditional transposon sequencing strategies used in mycobacteria. To create a resource for determining the function of genes, we generated a pooled random barcode transposon-site sequencing (RB-TnSeq) library in *Mycobacterium tuberculosis* (Mtb). A unique twenty-nucleotide barcode in the transposon allows for rapid, high-throughput genetic screening without the laborious protocol of standard bacterial TnSeq screens. We performed 95 RB-TnSeq screens on an array of carbon sources, nitrogen sources, stressors, and antibiotics. Using the resulting dataset, we examined phenotypes of PE/PPE genes, a mycobacterial gene family whose function has long been elusive, uncovering 187 novel phenotypes across 37 genes in this family. We propose a pathway for lactate utilization in which the ESX-5 type VII secretion system exports PPE3, facilitating the import of D- and L-lactate into the bacterial cell. Notably, we identify a candidate D-lactate dehydrogenase that may mediate this metabolic capability. Additionally, we find that the proton-pumping NADH dehydrogenase Nuo is required for utilization of propionate, highlighting the metabolic flexibility of Mtb. Lastly, we characterize a novel mutant that confers resistance to the new tuberculosis antibiotic pretomanid. Results from these genetic screens will facilitate the development of additional new hypotheses about the function of uncharacterized genes and will expand our knowledge of Mtb metabolism and resistance to stress.

## Introduction

*Mycobacterium tuberculosis* (Mtb) is a bacterium that infects the lungs and is the causative agent of tuberculosis, which is estimated to infect a quarter of the global population [1]. The success of Mtb as a human pathogen may be attributed to many unique features of mycobacterial physiology. Mtb resides in the phagosome of macrophages, which activate an arsenal of host defenses to attempt to eradicate the infection. For instance, the phagosome is acidified by the vacuolar-H(+)-ATPase, NOS2 catalyzes production of nitric oxide, and the NADPH oxidase generates a respiratory burst of superoxide. Although these immune processes are successful at killing many pathogens, Mtb can persist and establish a chronic infection.

To grow inside the host, Mtb must assimilate host-derived macromolecules including fatty acids, amino acids, and carbohydrates to use as carbon and nitrogen sources. A large portion of the Mtb genome is seemingly dedicated to lipid metabolism [2].

Consistent with this, mutants lacking the fatty acid transporter *mce1* or the cholesterol importer *mce4* exhibit attenuated virulence in murine models of infection [3,4], supporting the idea that fatty acids and cholesterol are important *in vivo* carbon sources. However, the range of carbon sources used by Mtb during infection remains unresolved. For instance, Mtb grows *in vitro* on L-lactate [5,6], a metabolite produced from the metabolism of carbohydrates. L-lactate is present in guinea pig granulomas [7] and is produced by immune cells during Mtb infection due to the Warburg effect, the metabolic shift from oxidative phosphorylation to aerobic glycolysis [8–11]. An L-lactate dehydrogenase mutant in Mtb is attenuated for infection in human macrophages [5] and is under positive selection in Mtb clinical strains [6], suggesting that molecules other than lipids may act as carbon sources *in vivo*. Interestingly, there is little research into whether Mtb can use the stereoisomer D-lactate as a carbon source.

This metabolic flexibility of Mtb reflects its ability to survive hostile conditions, which also extends to antibiotic exposure. Mtb is both intrinsically resistant to many antibiotics and acquires *de novo* resistance to clinically used antibiotics. As a result, antibiotic therapy for TB requires combinatorial therapy that presents significant side effects. Depending on the bacterial strain, treatment can range from three to nine months in duration [12], resulting in patient non-adherence and the rise of multidrug-resistant (MDR) and extensively drug-resistant (XDR) strains. Drug-susceptible TB regimens include antibiotics such as isoniazid, rifampicin, pyrazinamide, and ethambutol. In response to the rise of MDR and XDR Mtb strains, the WHO updated their guidelines in 2022 to include the BPaLM regimen [13], consisting of bedaquiline, pretomanid, linezolid, and moxifloxacin. Pretomanid, also referred to as PA-824, is a nitroimidazole that inhibits the synthesis of cell envelope ketomycolates [14]. The full spectrum of mutations that confer antibiotic resistance or susceptibility is yet to be determined for most antibiotics, particularly for newer drugs like pretomanid. Thus, the ability to predict what mutations may arise could be a clinically powerful tool.

Mtb encodes ∼4,000 genes, a similar size to the model bacterium *Escherichia coli* [15], yet many genes remain unannotated as “hypothetical proteins” or annotated only by homology with other bacteria [16]. The requirement for high-containment facilities and the slow growth of Mtb (21 days to grow a colony on solid agar) complicate laboratory experiments. In bacteria, genetic screens are often accomplished using transposon insertion sequencing (TnSeq), which couples pooled transposon insertion mutagenesis with next-generation sequencing [17–20]. Transposon-mutant libraries are exposed to a condition of interest, and the enrichment or depletion of transposon mutants are used to infer gene function. TnSeq screens in Mtb have revealed essential genes in broth [21–23] and genes important for virulence in various mouse models [24–26]. TnSeq screens have also been conducted in biologically-relevant conditions, some of which include acid and oxidative stress [27], tuberculosis antibiotics [28], hypoxia [29], and growth on cholesterol [22]. A major limitation of TnSeq methods are the long and laborious protocols for preparing the sequencing libraries.

To address the sample preparation limitations of TnSeq in bacteria, random barcode transposon-site sequencing (RB-TnSeq) was developed and has since been applied to diverse environmental and commensal bacterial species [30–32]. In RB-TnSeq, each transposon contains a unique twenty nucleotide barcode. One round of traditional TnSeq is performed on the initial library to identify the genetic location of each transposon insertion and its associated barcode. Then, barcode abundance can simply be quantified by PCR and deep sequencing of DNA barcodes (BarSeq), allowing for rapid, high-throughput screening.

In this study, we generated an RB-TnSeq library in Mtb and systematically profiled genetic requirements across a diverse panel of carbon sources, nitrogen sources, immune-relevant stressors, and antibiotics. The resulting dataset enabled functional insights into the enigmatic PE/PPE family genes, a mycobacterial gene family named after their conserved proline and glutamate motifs. These findings reveal a lactate utilization pathway in which ESX-5 secretes PPE3, facilitating the uptake of lactate, where it is converted into pyruvate by stereospecific lactate dehydrogenases.

Additionally, we find that the NADH dehydrogenase Nuo is essential for propionate utilization, offering a metabolic insight into why Mtb encodes three distinct NADH dehydrogenases. We also identify a putative operon involved in resistance to pretomanid, a recently approved antibiotic for drug resistant Mtb strains. These high-throughput genetic screens uncover novel aspects of Mtb biology. Elucidating molecular mechanisms behind drug susceptibility and resistance, metabolic flexibility, and resistance against immune stressors will be imperative for developing new therapeutics to combat the global tuberculosis epidemic.

## Results

### High-throughput fitness data with Mtb RB-TnSeq library

To create an RB-TnSeq library in Mtb, we ligated the temperature-sensitive mycobacteriophage vector phAE159 with pMtb_NN1, a plasmid containing a transposase under the T6 mycobacterial promoter, a *Himar1* mariner transposon with unique twenty-nucleotide barcodes, and a kanamycin resistance cassette (S1 Figure). The resulting phagemid was electroporated into *Mycobacterium smegmatis* and incubated at 30°C for lytic phage production. Subsequent transduction of the Mtb strain H37Rv with the mycobacteriophage at 37°C, the non-lytic temperature, yielded an RB-TnSeq library with ∼60,000 unique mutants across 2,888 genes. As TnSeq libraries in Mtb typically contain mutations in ∼3,300 genes [21,22], this RB-TnSeq library is not fully saturated. Generating the Mtb RB-TnSeq library presented considerable difficulty, largely due to the loss of barcode diversity at each step in the protocol. After much optimization, we proceeded with conducting genetic screens with the ∼60,000 barcode library.

To capitalize on the high-throughput nature of RB-TnSeq, we sought to conduct genetic screens across a chemical library of carbon sources, nitrogen sources, antibiotics, and stressor compounds. First, we tested if chemical library compounds met the growth parameters to conduct an RB-TnSeq screen. Carbon and nitrogen sources were dissolved in a modified Sauton’s minimal media, such that each carbon or nitrogen source was the predominant nutrient. A carbon or nitrogen source compound was used for RB-TnSeq screening if a minimal threshold of ∼3 doublings on the nutrient was observed (S1 Table and S2 Table). For antibiotics and stressors, IC50s were measured and an RB-TnSeq screen was then performed if the compound was inhibitory towards Mtb in the range of concentrations measured (S3 Table). In total, we tested 100 carbon sources, including sugars, nucleotides, amino acids, and lipids, 43 nitrogen sources, including amino acids and nucleotides, and 120 stressors and antibiotics.

After RB-TnSeq cultures reached saturation on their given compound, cultures were pelleted for genomic DNA extraction, followed by PCR amplification, and deep sequencing of DNA barcodes (BarSeq). The number of barcodes at the end of the experiment was compared to the number of barcodes at the beginning of the experiment, which is represented by the log2 fold change or “BarSeq fitness.” Mutants in genes that confer a growth advantage have a positive BarSeq fitness, while mutants in genes that confer a growth disadvantage have a negative BarSeq fitness (Figure 1A). A t-like statistic was computed that accounts for the consistency of all transposon mutants in a given gene [30,31]. We considered a gene to have a statistically significant phenotype in a given condition if |gene fitness| >0.5 and |t|>3.

**Figure 1.**
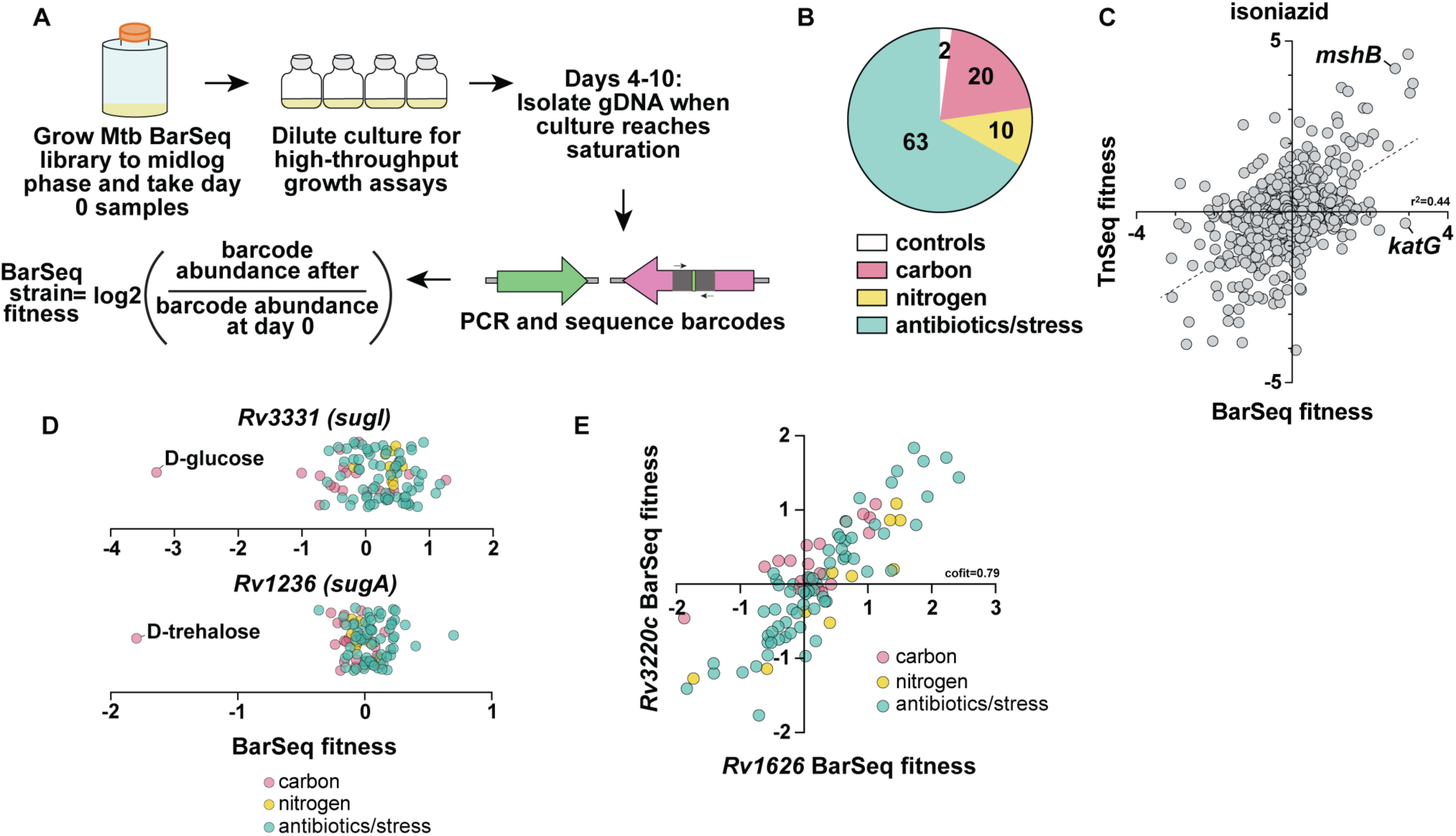
Overview of RB-TnSeq screens. (A) Schematic for RB-TnSeq screen experimental set-up. (B) Pie chart depicting the number of RB-TnSeq screens completed for each category. (C) Previously published TnSeq fitness with 27ng/mL isoniazid [28] plotted against BarSeq fitness with 25ng/mL isoniazid. Dotted line represents linear correlation between statistically significant hits from RB-TnSeq with TnSeq. (D) BarSeq fitness (log2 fold change) for *Rv3331 (sugI)* and *Rv1236 (sugA).* (E) Cofitness data for *Rv3220c* and *Rv1626*.

Of the 100 carbon sources we tested, 20 compounds met the growth parameters for BarSeq (Figure 1B). 9 of the successful carbon screens were lipids, which aligns with the well-characterized ability of Mtb to utilize lipids as carbon sources. However, we also conducted screens on 2 sugars (D-glucose and trehalose), glycerol, 4 amino acids (aspartic acid, glutamic acid, asparagine, and casamino acids), and cellular metabolites such as lactate, pyruvate, and malic acid. We did not observe growth on many of the sugars in the chemical library, such as D-fructose, D-xylose, sucrose, D-raffinose, and L-rhamnose. Interestingly, most amino acids did not support growth as a singular carbon source. Of the 43 nitrogen sources we tested, 10 compounds met the growth parameters for BarSeq, comprised of 8 amino acids, urea, and ammonium chloride. Nucleotides, such as uridine, thymidine, inosine, cytidine, thymine, and cytosine, did not result in robust growth as a singular carbon source or nitrogen source. However, these *in vitro* studies on minimal media do not exclude the possibility that these compounds may support survival of Mtb *in vivo*. Lastly, of the 120 stressors and antibiotics, 63 conditions were used for RB-TnSeq. Some stressors were conducted with multiple concentrations, resulting in 59 unique stressor and antibiotic compounds. Stressors included compounds such as hydrogen peroxide, nitric oxide, metals, and acidic pH. Antibiotics ranged from those not clinically used for TB (e.g. sisomicin, ciprofloxacin, doxycycline), ones traditionally used for TB treatment (e.g. rifampicin, isoniazid, pyrazinamide, and ethambutol), as well as new TB drugs (e.g. pretomanid, delamanid, and bedaquiline). We also performed controls alongside all screens – either a “7H9 no stress control”, our standard media for mycobacterial growth, or “7H9 with 1% DMSO” as a control for compounds dissolved in DMSO.

In total, we conducted 212 successful RB-TnSeq screens representing 95 unique experimental conditions in biological replicates (Figure 1B). The “7H9 no stress control” was performed with each experimental batch and was therefore repeated numerous times. From these data, we calculated 7524 total statistically significant hits in 850 unique genes. It is important to note that for high-throughput screening, we conducted two replicates for each condition. However, additional replicates can be conducted in the future for generating additional high-confidence hits for any condition of interest. Of the 95 conditions, only 6 have existing TnSeq datasets published in the literature to our knowledge. These include cholesterol [22], acidic pH (pH 4.5) [25], and treatment with the antibiotics isoniazid, rifampicin, ethambutol, and meropenem [28]. When comparing our RB-TnSeq data to these previously published TnSeq results, we observe strong concordance in gene-level phenotypes. Most genes with the strongest phenotypes are shared across the previously published TnSeq and our RB-TnSeq screens on isoniazid (Figure 1C), rifampicin, meropenem, and ethambutol (S2 Figure) [28]. For instance, *mshB (Rv1170)*, a mycothiol synthesis protein, confers resistance to isoniazid according to both TnSeq and RB-TnSeq, supporting previous findings connecting mycothiol synthesis with isoniazid resistance [33]. Mutants in *katG (Rv1098c)*, a catalase-peroxidase known for activating the isoniazid prodrug, was identified as a top hit for isoniazid resistance in the RB-TnSeq screen yet was not a hit in the published TnSeq screen (Figure 1C). Mutants in *pirG (Rv3810)*, an extracellular repetitive protein, are known to be susceptible to rifampicin [34], which was also a shared hit between TnSeq and RB-TnSeq (S2 Figure). In accordance with its annotation as a beta-lactamase, *blaC (Rv2068c)* was a shared hit between TnSeq and RB-TnSeq for exposure to meropenem (S2 Figure). The antibiotic concentration, time point, number of replicates, and differences in gene coverage in the mutant libraries may underlie differences between the TnSeq and RB-TnSeq.

Using our high-throughput fitness data, we first examined specific phenotypes, a metric in which a given gene only has a strong fitness defect (|fitness|>1 and |t|>5|) in one or a few conditions [31]. Specific phenotypes can be particularly useful in defining the function of a given gene. For example, *Rv3331*, annotated as a probable sugar-transport integral membrane protein *sugI*, was only attenuated for growth on D-glucose (Figure 1D), which supports a specific function as a sugar importer. Although *sugI* has not been shown to import D-glucose experimentally, it has been predicted by homology to transport monosaccharides [35]. In contrast, the *sugA (Rv1236)* operon, which is known to recycle the disaccharide trehalose [36], was specifically attenuated for growth on D-trehalose (Figure 1D). Our compilation of fitness data yielded 241 specific phenotypes for 169 unique genes (S4 Table).

We also evaluated cofitness, wherein multiple genes share the same fitness score across many conditions, suggesting the genes work together in a similar process or pathway [31]. Cofitness is calculated using the Pearson correlation of the fitness values between two genes [31]. We observed high cofitness values for 161 unique gene pairs, in which the cofitness score was greater than 0.8 (S5 Table). For instance, *Rv3220c* and *Rv1626* are putative members of a putative bacterial two-component system [37,38].

These two genes have a cofitness score 0.79, supporting existing biochemical data demonstrating a physical interaction (Figure 1E). In addition to their co-fitness, *Rv3220c* and *Rv1626* are both statistically significant hits in 16 different conditions including resistance to antibiotics like norfloxacin, ciprofloxacin, meropenem, linezolid, and isoniazid and hyper-susceptibility to pyrazinamide and para-aminosalicylic acid. These pleiotropic effects suggest this two-component system may have a global effect on the cell, supporting its previous association with lipid metabolism [38].

### Diverse phenotypes of PE and PPE genes

PE/PPE genes are a gene family unique to mycobacteria and are highly represented in Mtb, comprising 10% of coding capacity of the genome [2]. PE/PPE genes are defined by conserved proline (P) and glutamic acid (E) residues in the N-terminus. PE/PPE proteins can be secreted as heterodimers to the bacterial surface, where they are hypothesized to participate in nutrient acquisition and modulation of the immune response [39]. Given the significant expansion of PE/PPE genes in Mtb, many questions remain regarding the exact function of this unique gene family, as many members have yet to be characterized. Of the 168 PE and PPE genes encoded by Mtb, 154 of them were represented in the Mtb RB-TnSeq library. Our screening results demonstrate that PE/PPE genes display a wide array of fitness phenotypes across the screening conditions (Figure 2A). We calculated 184 statistically significant hits in 37 PE/PPE genes (S6 Table). Notably, many PPE/PE genes had zero to very mild fitness defects. This may support findings of PE/PPE proteins modulating the host immune system, which would not be represented in our *in vitro* screens. For example, *pe-pgrs33 (Rv1818c),* which has been reported to interact with TLR2 [40], had zero significant phenotypes on the conditions tested (Figure 2A). It is also possible that PE and PPE proteins may act redundantly, or be responsive to infection conditions, complicating the ability to observe strong phenotypes.

**Figure 2.**
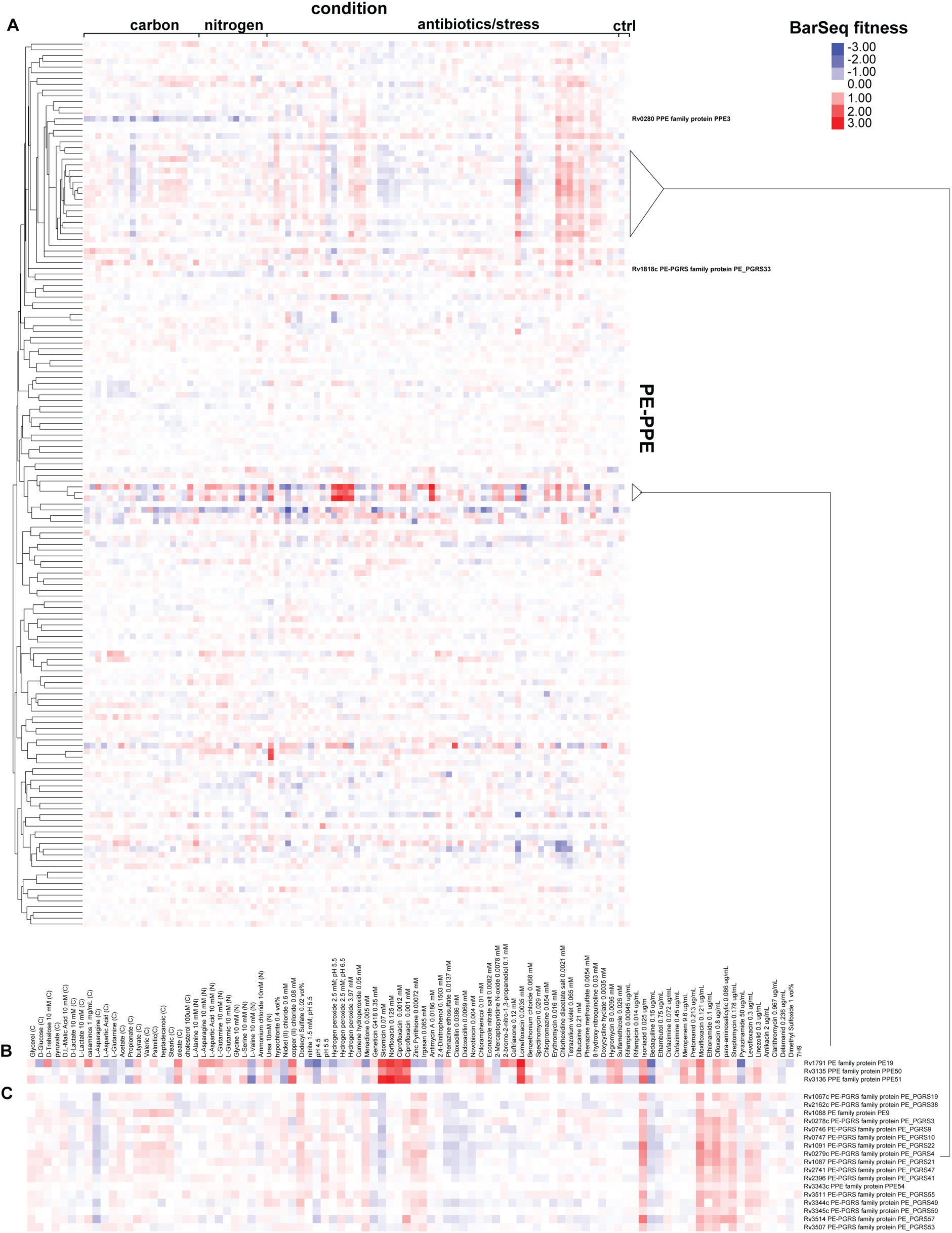
Heatmap of Mtb PE and PPE proteins. (A) Hierarchical clustering of BarSeq fitness values of 154 PE and PPE proteins represented in the RB-TnSeq library. (B) Gene cluster containing PE19, PPE50, and PPE51. (C) Gene cluster enriched in PE-PGRS genes with high cofitness values.

Hierarchical clustering revealed that some of the strongest phenotypes shown by the PE-PPE proteins were in PE19, PPE50, and PPE51 (Figure 2B). The clustering and high cofitness of these genes support previous findings that PPE51 interacts with PE19 [41]. PPE51 has been shown to be important for uptake of glycerol, glucose, maltose, lactose, and trehalose [41–43]. Our RB-TnSeq data additionally demonstrated mild growth defects for PPE51 on propionate, L-aspartic acid, and L-glutamine as carbon sources. In addition, we found that mutation of PE19, PPE50, or PPE51 resulted in resistance to sisomicin, norfloxacin, ciprofloxacin, lomefloxacin, isoniazid, and copper II chloride. Whether PPE50 interacts with PE19 and PPE51 or these phenotypes are due to a polar effect is yet to be experimentally determined.

Some of the strongest cofitness phenotypes we observed were between *pe9*, *ppe54*, and 14 different PE-PGRS genes (Figure 2C). PE-PGRS proteins are a subset of PE proteins that contain polymorphic GC-rich sequences (PGRS) [44]. Transposon mutants in this group of PE-PGRS genes display resistance to antibiotics, including isoniazid, moxifloxacin, ethioniamide, ofloxacin, para-aminosalicylic acid, streptomycin, levofloxacin, and linezolid. We hypothesize that this gene cluster also mediates import of these antibiotics into the cell or may mediate permeability of the cell envelope. High cofitness values suggests that these PE-PGRS proteins act in the same pathway. Whether they bind each other, or are important for each other’s secretion, or some alternative model is yet to be determined.

### PPE3 and ESX-5 are required for growth on diverse nutrient sources

Given previous work linking PE/PPE proteins to nutrient acquisition, we investigated whether specific PE/PPE genes are required for optimal growth on carbon or nitrogen sources. Notably, *ppe3* (*Rv0280*) mutants exhibited a growth defect across multiple carbon and nitrogen conditions (Figure 2A). *Ppe3* mutants showed reduced fitness when Mtb was grown on glycerol, D-glucose, D-lactate, L-lactate, propionate, and L-asparagine as sole carbon sources as well as on L-serine and L-asparagine as nitrogen sources (Figure 3A). Importantly, *ppe3* mutants were resistant to several antibiotics, including isoniazid, ofloxacin, and moxifloxacin, suggesting that loss of *ppe3* does not create a general permeability defect (Figure 3A). *Ppe3* has been shown to be upregulated in proteasome mutants and regulated by Zur, a zinc regulator [45,46]. To our knowledge, aside from transcriptional regulation, *ppe3* has not been experimentally characterized. To validate our RB-TnSeq results, we isolated a mutant with a transposon insertion in *ppe3* (*ppe3::tn*) from our arrayed transposon mutant library. This mutant grew normally in standard 7H9 broth culture (S3 Figure). RB-TnSeq indicated that *ppe3* mutants are unable to grow normally on either L- or D-lactate (Figure 3A). To validate these results, we tested whether the *ppe3::tn* mutant grows when L- or D-lactate is provided as the sole carbon source. We observed that whereas wild-type Mtb grows robustly on both D and L-lactate, the *ppe3::tn* mutant fails to grow to the same degree as WT on either isomer over the course of 12 days (Figures 3B and 3C). A complementation strain expressing *ppe3* under its native promoter also grew robustly on D- and L-lactate (Figures 3B and 3C). Because the loss of the lipid phthiocerol dimycocerosate (PDIM) results in a growth advantage in Mtb, we verified that the *ppe3* complemented strain and all strains in this study retain PDIM by assaying for the sensitivity to vancomycin in the presence of propionate [47] (S4 Figure). Thus, the PPE3 protein is required for utilization of both D- and L-lactate carbon sources.

**Figure 3.**
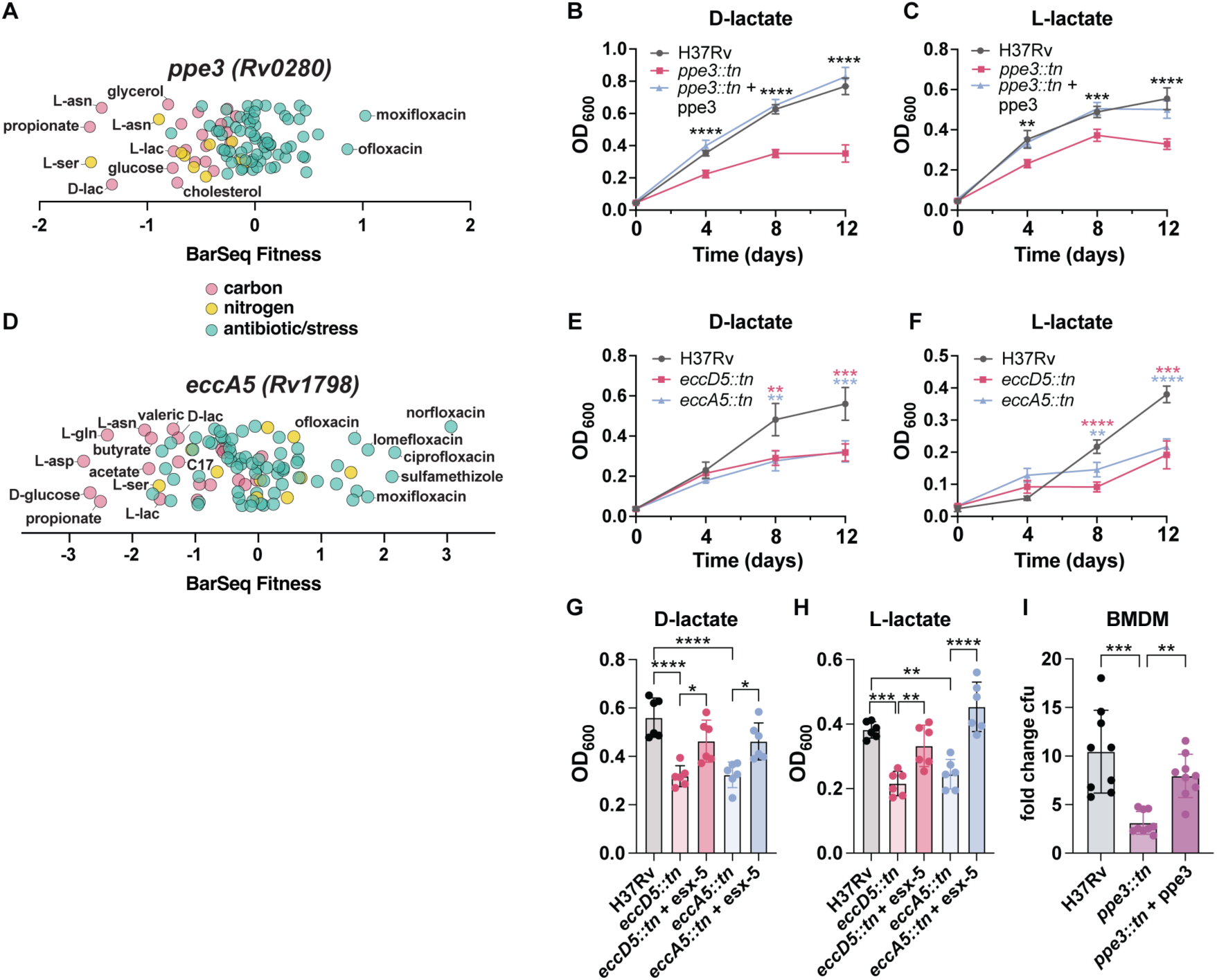
*ppe3* and *esx-5* are important for growth on nutrient sources. (A) BarSeq fitness of *ppe3.* (B) OD_600_ of WT, *rv0280::tn*, and complemented strain on 10mM D-lactate or (C) 10mM L-lactate over 12 days. (D) BarSeq fitness of *eccA5*. (E) OD^600^ of WT, *eccA5::tn*, or *eccD5::tn* on 10mM D-lactate or (F) 10mM L-lactate over 12 days. (G) Complementation of *eccD5::tn* and *eccA5::tn* in 10mM D-lactate or (H) 10mM L-lactate on day 12. (I) Fold-change CFU on day 4 from infection of mouse bone marrow-derived macrophages with H37Rv, *ppe3::tn*, and complemented strain. For B-H, data represent mean ± sd for *n*=6 replicates from two independent experiments. For (F), data represent mean ± sd for *n*=5 replicates from two independent experiments. p-Values were determined using two-way ANOVA for (B), (C), (E), and (F), and one-way ANOVA for (G) and (H) with Tukey’s multiple comparisons test. For (I), data represent mean ± sd for *n*=9 replicates from three independent experiments. p-Values were determined using Kruskal-Wallis test with Dunn’s multiple comparisons test (*p<0.05, **p<0.01, ***p<0.001, ****p<0.0001).

PPE and PE proteins have been found to be transported to the cell surface by ESX-5, one of the type VII secretion systems encoded by Mtb [48]. Thus, we sought to determine which nutrients require ESX-5 for their utilization. Mutation of the secretory component *eccA5 (Rv1798)* yields negative BarSeq fitness values for propionate, D-glucose, L-lactate, D-lactate, L-serine, acetate, butyrate, L-asparagine, L-glutamine, L-aspartate, valerate, and heptadecanoic acid (Figure 3D). These results agree with previous published literature demonstrating that ESX-5 is required for efficient utilization of fatty acids in *Mycobacterium marinum* [49]. To test whether ESX-5 is indeed required for growth on L- and D-lactate, we isolated Mtb strains with transposon insertions in two different genes of the *esx-5* locus, *eccA5::tn* and *eccD5::tn*. *EccA5* encodes the AAA+ ATPase, and *eccD5* encodes a putative channel protein of the ESX-5 secretion system. These *esx-5* mutants grow at the same rate as WT in standard growth medium (S3 Figure). Importantly, both mutants fail to grow robustly on both D- and L-lactate over the course of 12 days when compared to WT Mtb (Figures 3E and 3F).

Complementation of the ESX-5 operon restored the ability of *eccD5::tn* and *eccA5::tn* to grow on D- and L-lactate (Figures 3G and 3H).

Disruption of ESX-5 in Mtb is known to lead to a virulence defect in macrophages and mice [48]. Given the connection between ESX-5 and PE/PPE proteins and our observation that PPE3 is required for growth on various nutrients, we investigated whether PPE3 is important for growth in host cells. Enumeration of colony forming units (CFU) from a murine bone marrow-derived macrophage infection revealed that *ppe3::tn* is attenuated for growth in macrophages, which is rescued by the *ppe3* complement. (Figure 3I). Macrophage viability at this day four time point was comparable to uninfected cells and did not vary significantly between the WT, *ppe3::tn*, and complemented strains (S5 Figure).

### D-lactate as a potential carbon source for Mtb

The shared defect for *ppe3* and *esx5* mutants to grow on D- and L-lactate led us to further investigate lactate utilization in Mtb (Figure 4A). Growth on the distinct D and L isomers of lactate would require conversion into pyruvate by stereo-specific enzymes. The top RB-TnSeq hit for attenuated growth on L-lactate was *lldD2 (Rv1872c)*, which has previously been characterized as an L-lactate dehydrogenase [5] and is under positive evolutionary selection in Mtb clinical strains [6] (Figure 4A). As expected, a transposon mutant in *lldD2* was specifically attenuated for growth on L-lactate but not attenuated for growth on D-lactate or in standard 7H9 growth medium (Figures 4B and 4C, and S3 Figure). Expression of *lldD2* in this mutant strain restored growth on L-lactate (Figure 4E).

**Figure 4.**
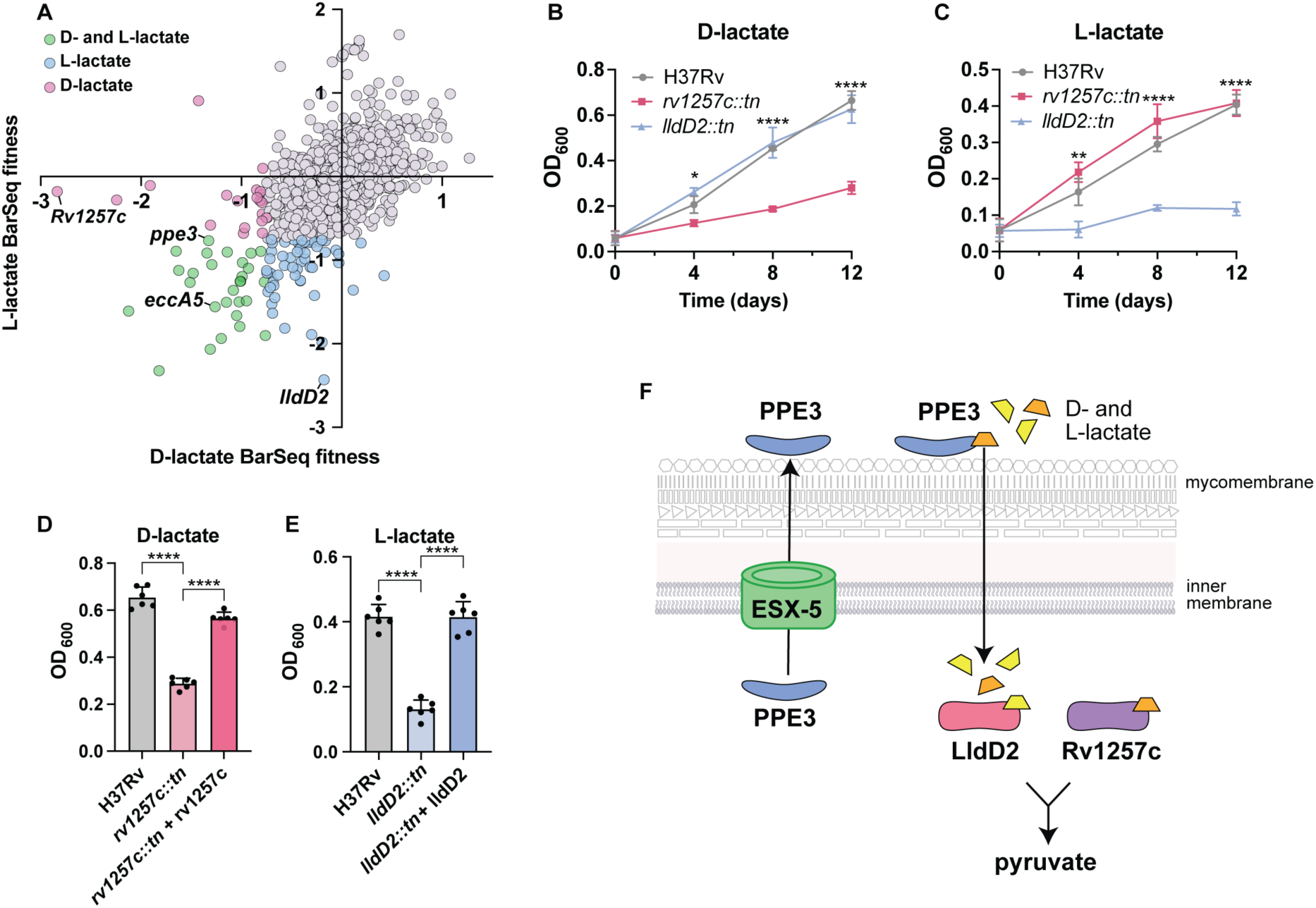
Mutants attenuated for growth on lactate. (A) L-lactate BarSeq fitness plotted against D-lactate BarSeq fitness. (B) OD^600^ of H37Rv, *rv1257c::tn,* and *lldD2::tn* on 10mM D-lactate and (C) 10mM L-lactate over 12 days. (D) Complementation of *rv1257c::tn* in 10mM D-lactate on day 12. (E) Complementation of *lldD2::tn* in 10mM L-lactate on day 12. (F) Model for lactate utilization in Mtb. For B-C, data represent mean ± sd for *n*=5 replicates from two independent experiments. For D-E, data represent mean ± sd for *n*=6 replicates from two independent experiments. p-Values were determined using two-way ANOVA for (B) and (C) and one-way ANOVA for (D) and (E) with Tukey’s multiple comparisons test (*p<0.05, **p<0.01, ***p<0.001, ****p<0.0001).

D-lactate can be produced from lipid and protein metabolism, but is largely produced as a product of carbohydrate metabolism. In macrophages, the glyoxalase system generates D-lactate from methylglyoxal, which is a reactive aldehyde produced as a side-product of glycolysis [50]. However, to our knowledge, D-lactate has yet to be implicated as a carbon source for Mtb. The top RB-TnSeq hit for growth on D-lactate was *Rv1257c*, a putative oxidoreductase (Figure 4A). Using a strain with a transposon insertion in *Rv1257c*, we confirmed that this mutant was specifically attenuated for growth on D-lactate (Figure 4B), as it grew to the same OD^600^ as WT on L-lactate and in standard 7H9 growth medium (Figures 4B and 4C, and S3 Figure). Complementation restored the ability for the *Rv1257c* mutant to grow on D-lactate (Figure 4D).

*Rv1257c* contains a potential FAD oxidase domain and shares 40.1% amino acid sequence identity with *E. coli* glycolate oxidase subunit D (*glcD*), which is a member of a protein complex that oxidizes glycolate to glyoxylate. *E. coli* glycolate oxidase uses D-lactate as a substrate at similar kinetics to glycolate [51]. Thus, we predict that *Rv1257c* acts as a D-lactate dehydrogenase in Mtb. Thus, RB-TnSeq has allowed us to propose a model where ESX-5 secretes PPE3 or other factors required for the import of D- and L-lactate. After D and L-lactate are imported into the cell, each isomer is converted to pyruvate by stereospecific lactate dehydrogenases (*lldD2* or *Rv1257c*) to enter central carbon metabolism (Figure 4F).

### The *nuo* operon is required for growth on propionate

Lipids are a prominent carbon source for Mtb during infection, so we conducted RB-TnSeq screens on a variety of fatty acid carbon chain lengths, ranging from acetate (C2) to stearate (C18). Metabolism of branched-chain fatty acids, odd-chain fatty acids, and cholesterol results in the production of the three-carbon molecule propionyl-CoA.

Propionyl-CoA is oxidized to pyruvate by the methylcitrate cycle through the action of methylcitrate synthase (*Rv1131*, *prpC),* methylcitrate dehydratase (*Rv1130, prpD*), and methyl-isocitrate lyase (*Rv1129c*, *icl1*) [52] (Figure 5B). Accordingly, top hits that conferred susceptibility to growth on propionate included *prpC*, *prpD*, and their regulator, *prpR* (Figure 5A).

**Figure 5.**
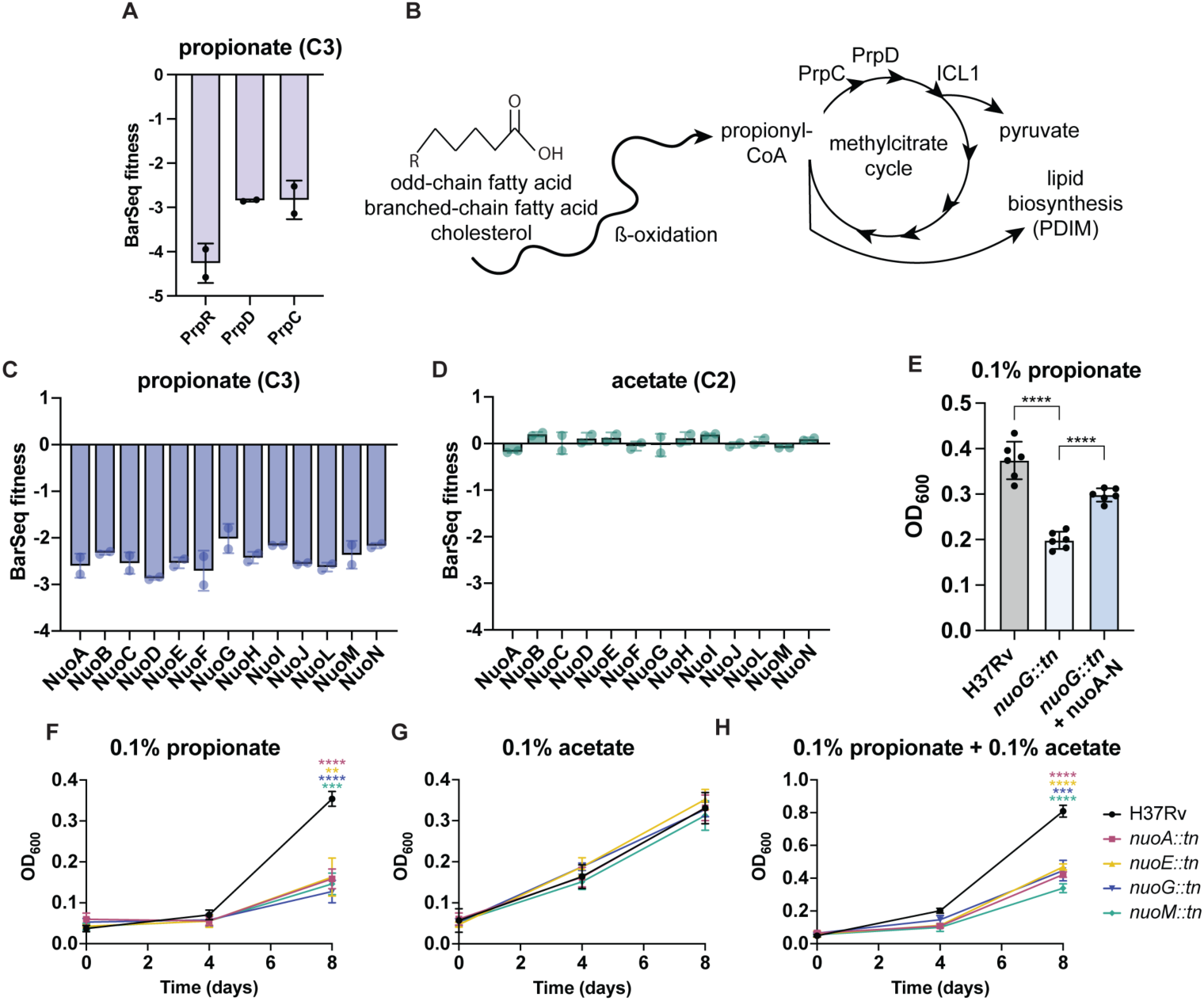
Nuo mutants are attenuated on propionate. (A) BarSeq fitness of methylcitrate cycle genes for growth propionate. (B) Schematic of propionate detoxification by the methylcitrate cycle or shuttling into lipid biosynthesis. (C) BarSeq fitness of the *nuo* operon for growth on propionate and (D) acetate. (E) OD^600^ of H37Rv, *nuoG::tn,* and complemented strain in 0.1% propionate on day 8. Data represent mean ± sd for *n*=6 replicates from two independent experiments. (F) OD^600^ of WT and *nuo* transposon mutants for growth on 0.1% propionate (G) 0.1% acetate, and (H) 0.1% propionate and 0.1% acetate on days 0, 4, and 8. Data represent mean ± sd for *n*=4 replicates from two independent experiments. p-Values were determined using one-way ANOVA for (E) and two-way ANOVA for (F), (G), and (H) with Tukey’s multiple comparisons test (*p<0.05, **p<0.01, ***p<0.001, ****p<0.0001).

Another top hit for growth on propionate was the 14-subunit NADH dehydrogenase complex encoded by the *nuo* operon (*nuoA* (*Rv3145)-nuoN (Rv3158*)) (Figure 5C), which was not a hit for growth on the two-carbon fatty acid acetate (Figure 5D). Mtb possesses three NADH dehydrogenases; *nuo, ndh* and *ndhA*. Nuo is a classical type I NADH dehydrogenase that oxidizes NADH to NAD+ and contributes to the proton motive force that drives ATP production by translocating protons to the outer membrane space. *Ndh* (*Rv1854c*) and *ndhA* (*Rv0392c*) differ from *nuo* and are classified as type II NADH dehydrogenases, consisting of one subunit that lacks proton-pumping capabilities.

Using mutants with transposon insertions in *nuoA*, *nuoE*, *nuoG*, or *nuoM*, we confirmed that *nuo* mutants are attenuated for growth on 0.1% propionate as the sole carbon source by measuring OD^600^ over the course of 8 days (Figure 5E and 5F).

Complementation of *nuoG::tn* with the *nuo* operon partially restored growth on propionate (Figure 5E). As predicted by the RB-TnSeq data, the *nuo* mutants grew to the same degree as WT on 0.1% acetate after 8 days in culture (Figure 5G). Due to propionate toxicity, bacteria that cannot metabolize propionate, such as methylcitrate cycle mutants, fail to grow in the presence of propionate and acetate [52]. We therefore tested for defects of the *nuo* mutants on a combination of 0.1% acetate and 0.1% propionate. We observed that the growth of *nuo* mutants on this mixed carbon source was attenuated compared to WT and mimicked the growth on 0.1% acetate alone (Figure 5H). Given that the *nuo* mutants can grow on acetate in the presence of propionate, we suspect that propionate is detoxified in the *nuo* mutants but cannot be used as a carbon source to its full potential. Although *nuoG* is a known virulence factor and *nuoG* mutants are attenuated in mouse infection models [53,54], this report is the first to identify a growth defect on propionate, which may contribute to the previously observed virulence attenuation.

### Mutants in *Rv1634* confer resistance to pretomanid

In addition to nutrients, we conducted RB-TnSeq screens to find genes that confer susceptibility and resistance to antibiotics and stressors. Pretomanid is a relatively new TB therapy that is included in new regimens for drug-resistant tuberculosis. It is administered as a prodrug that requires activation by deazaflavin-dependent nitroreductase (*Ddn*), an enzyme involved in the redox cycling of the cofactor F420 [55]. Mutations that confer resistance to pretomanid have largely been found in the genes that are responsible for activating pretomanid prodrug. These include *ddn*, F420 synthesis genes (*fbiA-D*), and *fgd1*, which reduces the F420 cofactor [56–59]. In alignment with these previous findings, top hits for pretomanid-resistant mutants in the RB-TnSeq screen included *ddn*, the *fbi* operon, and *fgd1* (Figure 6A).

**Figure 6.**
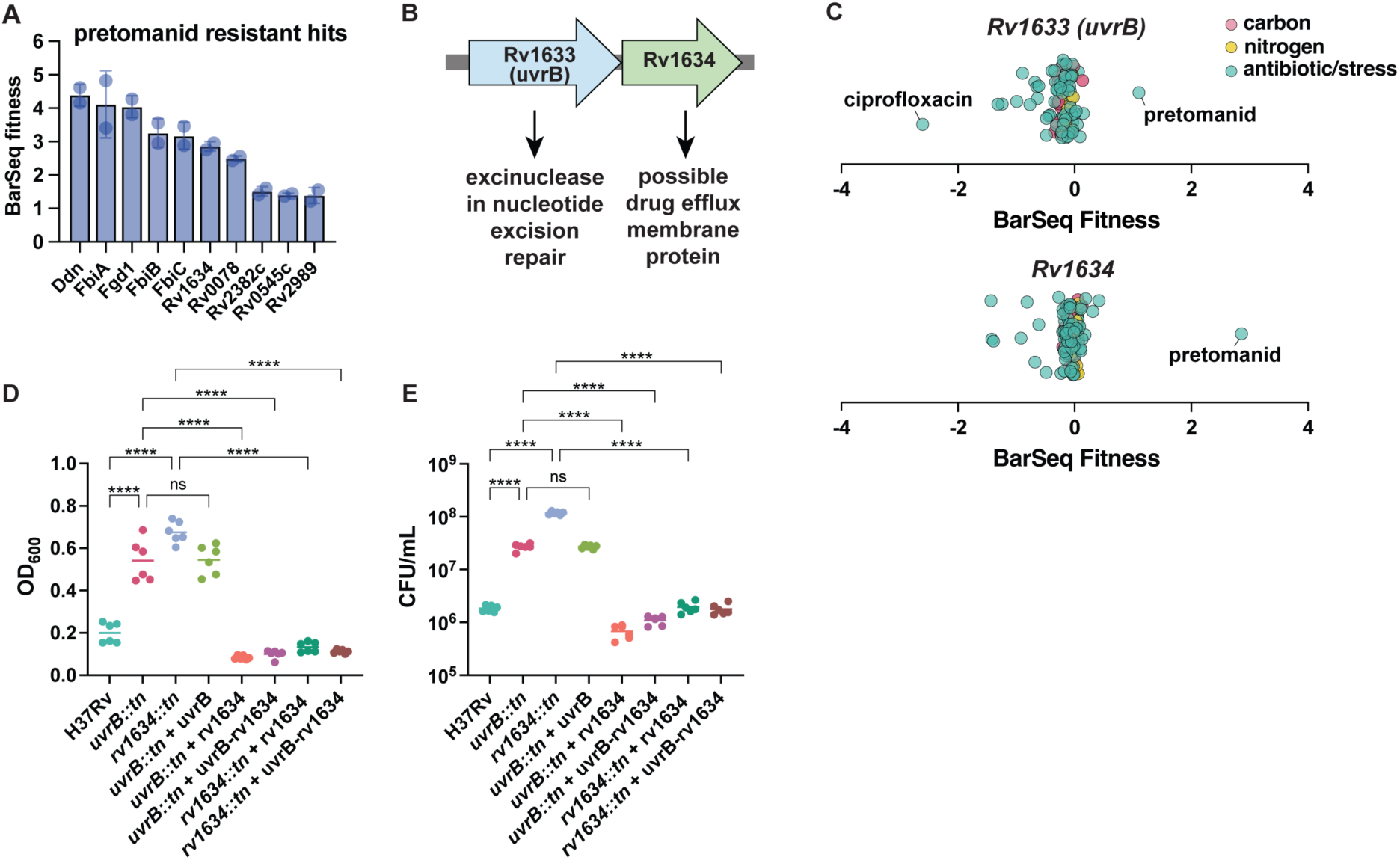
*Rv1634* confers resistance to pretomanid. (A) Top 10 BarSeq hits for pretomanid resistance. (B) *uvrB* and *Rv1634* (possible drug efflux membrane protein) are hypothesized to be in an operon. (C) BarSeq fitness of all RB-TnSeq screens for *uvrB* and *Rv1634*. (D) OD^600^ or (E) CFU enumeration of WT, *uvrB::tn*, and *rv1634::tn* and complemented strains after exposure to 0.45 µg/mL pretomanid for 5 days. Data represent mean ± sd for *n*=6 replicates from two independent experiments. p-Values were determined using one-way ANOVA with Tukey’s multiple comparisons test (*p<0.05, **p<0.01, *** p<0.001,****p<0.0001).

Besides the genes involved in prodrug activation, we find that *Rv1634,* encoding a putative drug efflux transporter, is the strongest hit conferring resistance to pretomanid (Figure 6A). *Rv1634* is operonic with *uvrB (Rv1633)*, an excinuclease in nucleotide excision repair that removes bulky lesions from DNA (Figure 6B). The RB-TnSeq data suggest that *uvrB* and *Rv1634* specifically confer resistance to pretomanid and not any other antibiotics tested in our screens (Figure 6C). Transposon mutants in *uvrB* and *Rv1634* (*uvrB::tn and rv1634::tn*) showed increased survival in the presence of pretomanid as measured by OD600 (Figure 6D) or when plating for CFU after 5 days of antibiotic exposure (Figure 6E), while untreated strains grew to the same degree as WT (S6 Figure). The CFU data demonstrate that *uvrB::tn* and *rv1634::tn* display approximately a 15-fold and 65-fold increase in pretomanid resistance respectively.

Complementation of *uvrB::tn* with *uvrB* alone failed to restore growth to WT levels in the presence of pretomanid (Figure 6D and E). However, complementation of *uvrB::tn* with *Rv1634* or *uvrB-Rv1634* successfully restored the susceptibility to pretomanid (Figure 6E). Thus, it appears pretomanid resistance is driven by *Rv1634.* It is unexpected that a loss-of-function mutant in a putative drug efflux protein drives susceptibility to an antibiotic. One potential explanation is that pretomanid enters the cell through this drug efflux membrane protein, but more experiments are needed to test this hypothesis.

Interestingly, *Rv1634* may confer resistance to pretomanid outside of the conventional *ddn*, *fgd1*, F420 redox recycling pathway typically associated with pretomanid-resistant mutants discovered thus far.

## Discussion

We performed 212 successful RB-TnSeq screens representing 95 conditions on a range of carbon sources, nitrogen sources, stressors, and antibiotics related to Mtb physiology. This high-throughput data allows generation of hypotheses about gene functions by applying specific phenotypes and cofitness measurements. From our screening data, we dissect a lactate utilization pathway whereby ESX-5 secretes PPE3, which uptakes D and L-lactate into the bacterial cell. Lactate is then converted to pyruvate by stereospecific lactate dehydrogenases and enters central carbon metabolism. We also show that the NADH dehydrogenase encoded by the *nuo* operon is essential for growth on propionate. Lastly, we uncover a novel gene conferring resistance to pretomanid, *Rv1634*, a putative drug efflux pump.

A major limitation of our RB-TnSeq screens is that the library is unsaturated. Maintaining large barcode diversity throughout the library construction process presented an enormous challenge. The RB-TnSeq library technically contains a large amount of unique barcoded transposon insertions, but many of these barcodes are not usable because they map to multiple insertion sites. Even if the library were saturated, we would lack representation of essential genes, which comprise ∼15-20% of the genome. To address this caveat of transposon libraries, CRISPRi libraries have been developed in Mtb and screened against TB antibiotics [60]. Nonetheless, data from these high-throughput genetic screens will provide a useful resource for deepening our understanding of mycobacterial genetics.

Our observation that Mtb grows robustly on D-lactate in broth as the sole carbon sources reveals a potential for D-lactate to be utilized during infection. D-lactate dehydrogenases have long been elusive in Mtb. To our knowledge, the sole enzyme with D-lactate dehydrogenase activity to have been characterized in Mtb is the flavohemoglobin *FHb* (*Rv0385*), which was shown biochemically to oxidize D-lactate to pyruvate by the action of FAD and heme cofactors [61]. A follow-up study on *FHb* revealed an additional biochemical function as a disulfide oxidoreductase [62]. In our dataset, *Rv0385* did not have significant BarSeq fitness defect for growth on D-lactate. Based on sequence homology to *E. coli*, we predict that *Rv1257c* may be the predominant D-lactate dehydrogenase in Mtb for utilization of D-lactate as a carbon source. For utilization of L-lactate, our findings match previous data that *lldD2* rather than *lldD1* is the primary L-lactate dehydrogenase in Mtb [5]. Interestingly, our dataset reveals that *lldD1* may be important for cholesterol utilization.

An interesting question is why Mtb encodes three different NADH dehydrogenases: *nuo*, *ndh*, and *ndhA*. Until recently, *ndh,* a non-proton pumping NADH dehydrogenase, was thought to be an essential gene because attempts at knocking out *ndh* were unsuccessful [63], and *ndh* was not represented in TnSeq libraries, including our RB-TnSeq library. However, *ndh* knockouts have now been generated in the CDC1551 strain [54] and in H37Rv in fatty-acid free media [64]. Transposon insertions have been found in the other non-proton pumping NADH dehydrogenase *ndhA* [21], although questions remain as to the exact roles of these three NADH dehydrogenases. Notably, *ndhA* was not a BarSeq hit for any screen conducted thus far. We hypothesize that Nuo is important when the cell is under certain types of metabolic stress because its proton-pumping capabilities allow for energy conservation due to the contribution to the proton motive force. It is also possible that proton pumping may help modulate the cytoplasmic pH. It is well-appreciated that Mtb is a metabolically flexible bacterium, and these data highlight how the apparent redundancy of three NADH dehydrogenases allows for growth on different carbon sources. It would be interesting to add radiolabeled propionate to bacterial cultures and track the fate of the radiolabeled carbon in the *nuo* transposon mutants.

The last phenotype we followed up on was concerning the TB antibiotic pretomanid, which is included in new regimens for drug-resistant TB. We found that mutants in *Rv1634* confer resistance to pretomanid, which we hypothesize may act as a stepping-stone to develop even higher resistance *in vivo*. *Rv1634* encodes a putative drug efflux membrane protein. This presents a counter-intuitive situation. Typically, a loss-of-function mutant in a drug efflux protein drives susceptibility to an antibiotic. However, we observe that a mutant in *Rv1634* drives resistance to pretomanid. We predict that pretomanid may enter the cell through *Rv1634.* Future experiments measuring intracellular pretomanid levels or conducting suppressor screens would be enlightening in determining the mechanism of *Rv1634*-driven pretomanid resistance. A deeper understanding of the functions of previously uncharacterized genes in Mtb will be imperative in the development of novel TB antibiotic therapies.

## Materials & Methods

### Bacterial strains and culture

The Mtb strain H37Rv was used for generation of the BarSeq library and for all subsequent experiments. Transposon mutants (*ppe3::tn, rv1257c::tn*, *lldD2::tn*, *eccA5::tn, eccD5::tn, nuoA::tn*, *nuoE::tn*, *nuoG::tn*, *nuoM::tn, uvrB::tn*, *rv1634::tn*) were picked from an arrayed transposon library generated at the Broad Institute. For the ESX-5 complements, pMV-ESX-5 encoding the ESX-5 operon was used as described previously [65]. For the *nuo* complement, the entire *nuo* operon and its upstream promoter was cloned into pMV306. The remaining complemented strains were generated by Gibson assembly cloning of the gene coding region plus 1000bp upstream into pMV306. For all experiments, Mtb was grown to midlog phase in Middlebrook 7H9 (Difco BD) liquid medium supplemented with 10% albumin-dextrose-saline, 0.4% glycerol, and 0.05% Tween80. When plated, Mtb was grown on solid 7H10 agar supplemented with Middlebrook OADC (BD Difco) and 0.4% glycerol. For growth on carbon and nitrogen sources, Sauton’s media was prepared as previously described [66], except tyloxapol was substituted for Tween-80. For carbon source experiments in Sauton’s minimal medium, glycerol was omitted and replaced by the respective carbon source. For nitrogen source experiments in Sauton’s minimal media, asparagine and ammonium iron citrate were omitted and replaced by the respective nitrogen source.

### Generation of random barcode transposon-site sequencing library

#### PacI Digestion

10 µg of phAE159 cosmid and 10 µg of pMtb_NN1 were PacI digested overnight at 37°C followed by heat inactivation for 10 minutes at 75°C. phAE159 digestion was purified by addition of 1:10 volume of 3M sodium acetate and 2.5 volumes of 100% ethanol. Mixture was incubated at -20°C for 10 minutes. DNA was pelleted, washed with 70% ethanol, and air-dried for 5 minutes. Pellet was resuspended in 15 µL TE buffer. pMtb_NN1 digestion was purified by DNA cleanup and concentrator kit (Zymo Research) and eluted in 15 µL of TE buffer.

#### T4 Ligation

3 µg digested phAE159 cosmid and 1.5 µg digested pMtb_NN1 were incubated with 100 units of T4 Ligase, HC at 16°C for 4 hours. The reaction was ethanol precipitated as described above and resuspended in 20 µL TE buffer.

#### In vitro lambda packaging and E. coli transduction

10 µL of ligated phAE159 + pMtb_NN1 was added to 25 µL of MaxPlax Packaging extract and incubated at 30°C for 90 minutes. Another 25 µL of MaxPlax Packaging extract was added to reaction mixture and incubated at 30°C for another 90 minutes. 500 µL of PD buffer (10mM Tris-HCl pH 8.3, 100mM NaCl, and 10mM MgCl2) was added to stop the reaction followed by addition of 25 µL chloroform. The packaging reaction was then titered to estimate the efficiency of the reaction. Packaged lambda phage was incubated with Stbl3 *E. coli* at 37°C for 1.25 hours with no shaking and gentle vortexing every 15 minutes. Bacteria were then pelleted at 3,500 rpm for 5 minutes and resuspended in LB broth. Bacteria were plated on 24.5 cm^2^ LB + 50 µg/mL kanamycin plates to achieve 60,000 colony forming units per plate. A total of 1x10^6^ CFUs were plated. The next day, colonies were collected by scraping into LB broth and phagemid was purified by midi-prep.

#### Electroporation of BarSeq phagemid into M. smegmatis to generate mycobacteriophage

Electrocompetent *M. smegmatis* was prepared by washing 2 L of *M. smegmatis* grown to OD^600^=0.2 with ice cold 10% glycerol and resuspending in a final volume of 20 mL 10% ice cold glycerol. 400 µL of electrocompetent *M. smegmatis* and 200 ng of phagemid were added to 0.2 cm electroporation cuvettes. Cuvettes were incubated on ice for 10 minutes before electroporation pulse of 2.5 kV, 25 µF, and 1,000Ω. 2 mL of LB broth was immediately added to the electroporation cuvette, transferred to 15 mL conicals, followed by incubation at 37°C for 2 hours with no shaking. 8 mL of top agar (2mM CaCl2, 0.6% agarose, melted and cooled to 55°C) was added, and seeded onto two 7H10 plates. Plates were incubated at 30°C for 3 days for plaque formation. A total of 100 electroporations were conducted to make the RB-TnSeq library.

#### Mycobacteriophage collection and concentration

Phage was collected by adding 3mL MP buffer (50mM Tris pH 7.6, 150mM NaCl, 10 mM MgCl^2^, and 2mM CaCl^2^) to each 7H10 plate and rocked at 4°C overnight. The liquid on each plate was collected, pooled into 50 mL conicals, and centrifuged at 4,000 rpm for 20 minutes to pellet any bacteria. The supernatant was then passed through a 0.22 µm filter. To concentrate phage, a 20% PEG-8000/2.5M NaCl mixture was incubated with the phage at a 7.5mL PEG-NaCl to 30 mL phage ratio on ice for 2 hours. The phage was pelleted at 15,000 rpm for 30 minutes at 4°C. Phage was resuspended in MP buffer and subsequently titered.

#### Mycobacterium tuberculosis transduction

1L of Mtb was grown to OD^600^=0.8, washed two times with MP buffer, and resuspended in a final volume of 9 mL MP buffer. 1 mL of phage (concentration between 1x10^11^ and 1x10^12^ pfu/mL) was added to the bacteria and incubated at 37°C for 18 hours without shaking. Transduction was then washed two times and resuspended in 10 mL PBS+0.05% Tween80. The transduction was plated on 7H10+50µg/mL kanamycin + 0.05% Tween80 24 cm^2^ plates and incubated at 37°C for 21 days. Colonies were collected by scraping into 7H9 broth. Bacterial clumps were dissolved by sonication and light vortexing, and the RB-TnSeq library was aliquoted into cryovials for storage at - 80°C.

### RB-TnSeq fitness assays

Mtb RB-TnSeq library was grown from frozen stocks in 7H9+50µg/mL kanamycin in roller bottles for 5 days at 37°C. For day 0 barcode abundance, 5mL of culture was pelleted and stored at -80°C until genomic DNA extraction. For carbon source experiments, the culture was washed 2x with Sauton’s minimal media minus carbon (no glycerol). For nitrogen source experiments, the culture was washed 2x with Sauton’s minimal medium minus nitrogen (no asparagine and ammonium iron citrate). For antibiotic and stress experiments, the culture was washed 2x with 7H9. Cultures were started at OD^600^=0.05 in 10 mL in inkwells in duplicate and shaken at 37°C. For carbon and nitrogen sources, the cultures were pelleted for genomic DNA extraction when the culture reached saturation (OD^600^=0.8-1) or had at least doubled three times (OD^600^>0.4). For antibiotic and stress experiments, cultures were pelleted on day 5. For antibiotic and stress experiments that were more than 50% inhibitory (e.g. acidic pH, see S3 Table), cultures were outgrown in regular 7H9 and were pelleted when the culture reached midlog phase.

### Mtb genomic DNA extraction

10mL of culture was pelleted and frozen at -80°C. When thawed, pellet was resuspended in 440 µL RB buffer (25mM Tris-HCl pH 7.9, 10mM EDTA, and 50mM D- glucose) with 1mg/mL lysozyme and 0.2mg/mL RNase A and incubated at 37°C overnight. 100 µL 10% SDS and 50µL 10mg/mL proteinase K were added and incubated at 55°C for 30 minutes. 200 µL 5M NaCl was added followed by gentle mixing. 160 µL of pre-heated cetrimide saline solution (4.1g NaCl and 10g of cetrimide (hexadecyltrimethylammonium bromide) in 90 mL H^2^O) was added and incubated at 65°C for 10 minutes. 1 mL of chloroform:isoamyl alcohol (24:1) was added and inverted to mix. Mixture was centrifuged at 14,000 rpm for 10 minutes, and 800 µL of aqueous layer was transferred to a new tube. 560 µL (0.7x volume) isopropanol was added and mixed to precipitate DNA. Tubes were centrifuged at 14,000 rpm for 10 minutes, DNA pellet was washed with 700 µL 70% ethanol, and centrifuged at 14,000 rpm for 5 minutes. Pellets were air-dried and covered in 50µL water. DNA was stored at 4°C overnight to allow for DNA pellets to dissolve.

### Growth curves for carbon sources

Mtb strains were grown to midlog phase in 7H9, washed two times with Sauton’s medium minus carbon, and diluted to OD^600^=0.05 in 10mL in inkwells in media containing the respective carbon source (10mM sodium D-lactate, 10mM sodium L-lactate, 0.1% sodium propionate, 0.1% sodium acetate). Cultures were shaken at 37°C, and OD^600^ was measured on days indicated in figure legends. All carbon sources are from Sigma-Aldrich.

### Determination of antibiotic IC50s in 96 well-plates

Mtb strains were grown to midlog phase in 7H9 and washed two times with fresh 7H9. Mtb was added to 96 well-plates containing two-fold dilutions of antibiotics for a final starting OD^600^ of 0.05 in 100µL. Plates were incubated in a standing incubator at 37°C in a sealed Tupperware with a water-wet rag to prevent evaporation for 10 days. OD^600^ was measured using a SpectraMax M2 Microplate Reader. IC50s were determined by a nonlinear regression on GraphPad Prism.

### Determination of pretomanid resistance in culture

Mtb strains were grown to midlog phase in 7H9 and washed two times with fresh 7H9. Mtb was added to inkwell bottles containing 0.45 µg/mL pretomanid (Sigma-Aldrich) for a final starting OD^600^ of 0.05 in 10mL. Cultures were shaken at 37°C for 5 days. CFUs were determined by diluting bacteria into PBS+0.05% Tween-80 and plating serial dilutions on 7H10.

### Bone marrow-derived macrophage infections

Bone-marrow macrophages were collected from mice through a protocol approved by the UC Berkeley Institutional Animal Care and Use Committee (IACUC), protocol AUP- 2015-09-7979-3. Femurs from C57BL/6 mice were flushed and resulting cells were cultured in Dulbecco’s Modified Eagle Medium (DMEM) supplemented with 10% fetal bovine serum (FBS) and 10% supernatant from 3T3-M-CSF cells for 6 days. Two days before infection, BMDMs were seeded at a concentration of 50,000 cells per well in a 96-well plate. Bacteria were prepared by washing two times with PBS, sonicated, and spun at 500 rpm for 5 minutes to pellet clumps. Bacteria were diluted into DMEM supplemented with 5% FBS and 5% horse serum at a multiplicity of infection (MOI) of 1. Following a 4 hour phagocytosis period, infection medium was removed, and cells were washed with room temperature PBS before fresh medium was added. For CFU enumeration, medium was removed and cells were lysed in water with 0.5% Triton-X and incubated at 37°C for 10 minutes. Following the incubation, lysed cells were resuspended and serially diluted in PBS with 0.05% Tween-80. Dilutions were plated on 7H10 plates.

### Cell viability assay

The 96-well plate with bone marrow-derived macrophages was equilibrated to room temperature for 30 minutes. Following a PBS wash, 100µL of Cell Titer Glo 2.0 (ProMega) was added per well. The plate was incubated for 10 minutes at room temperature. Luminescence was measured in a white opaque 96 well-plate with a SpectraMax M2 Microplate Reader.

### Van-10-P assays

Van-10-P assays were conducted as described previously [47]. Briefly, Mtb strains were inoculated into 96-well plates at a starting OD^600^=0.005 in 7H9 with 0.1mM sodium propionate and 0.05% tyloxapol with or without 10µg/mL vancomycin (ThermoScientific). Plates were incubated for 10 days at 37°C at which the OD^600^ was measured using a SpectraMax M2 Microplate Reader. Van-10-P growth % was calculated as the ratio of the OD^600^ of vancomycin-treated wells over the untreated wells. Mtb Erdman lacking PDIM (PDIM-) was used as a control and described previously [67].

### Data analysis

TnSeq sequencing and data analysis, BarSeq, computation of gene fitness values, t scores, specific phenotypes, cofitness, and quality metrics were done as described previously [30]. Genes were considered to have significant BarSeq fitness if the |log2 fold change|>0.5 and |t-like statistic|>3. Hierarchical clustering of PE/PPE genes was computed in Gene Cluster 3.0 and visualized in Java TreeView. All other statistics were calculated in GraphPad Prism 10. For p-values, *p<0.05, **p<0.01, ***p<0.001, ****p<0.0001.

### Data availability

RB-TnSeq screening data are available on the public fitness browser (https://fit.genomics.lbl.gov/cgi-bin/org.cgi?orgId=MycoTube) or available on FigShare (https://figshare.com/s/f167c433d423d95920b9).

Reserved DOI: 10.6084/m9.figshare.30444569

## Acknowledgements

We thank Morgan Price for help with data analysis. We thank Edith Houben for the ESX-5 complementation plasmid. We thank members of the Stanley, Cox, and Vance labs for helpful discussions about this work.

## Supporting Information

**S1 Table. Carbon sources tested in this study**

**S2 Table. Nitrogen sources tested in this study**

**S3 Table. Antibiotics and stressors tested in this study**

**S4 Table. Specific phenotypes**

**S5 Table. Genes with high cofitness**

**S6 Table. PE/PPE genes with statistically significant BarSeq hits**

---

**S1 Figure.**
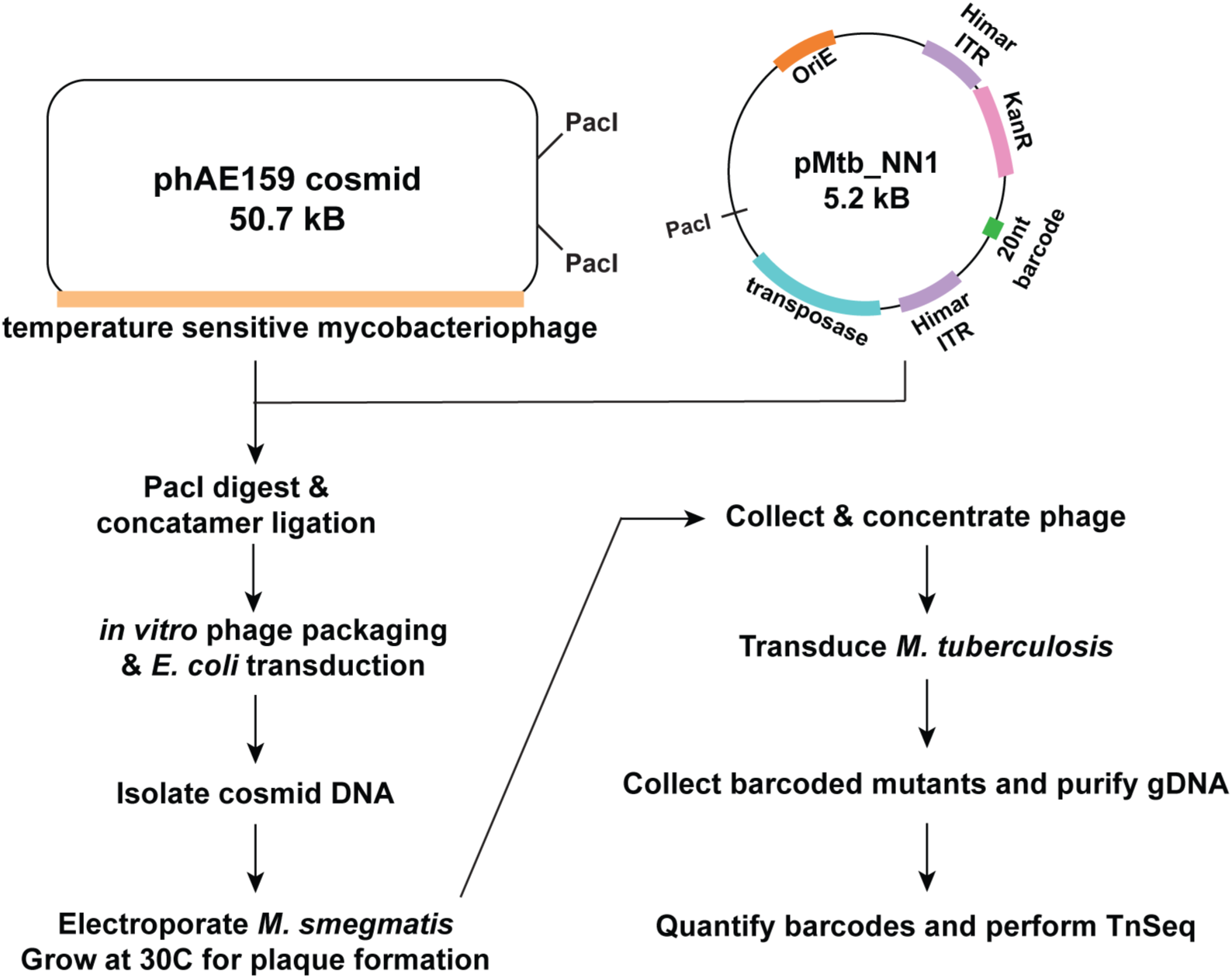
Protocol for construction of RB-TnSeq library in Mtb. Temperature-sensitive phAE159 cosmid was combined with pMtb_NN1 containing the transposase, *Himar1* mariner barcoded transposons, and a kanamycin resistant cassette via PacI digest and concatemer ligation. The ligated cosmid was packaged into lambda phage and transduced into *E. coli* to be midi-prepped. Barcoded phagemid was electroporated into *Mycobacterium smegmatis* and incubated at 30C for lytic phage plaque formation. Once the phage was collected and concentrated, it was transduced into Mtb and plates were incubated at 37C. After 21 days, colonies were scraped from plates and pooled to create the RB-TnSeq library.

**S2 Figure.**
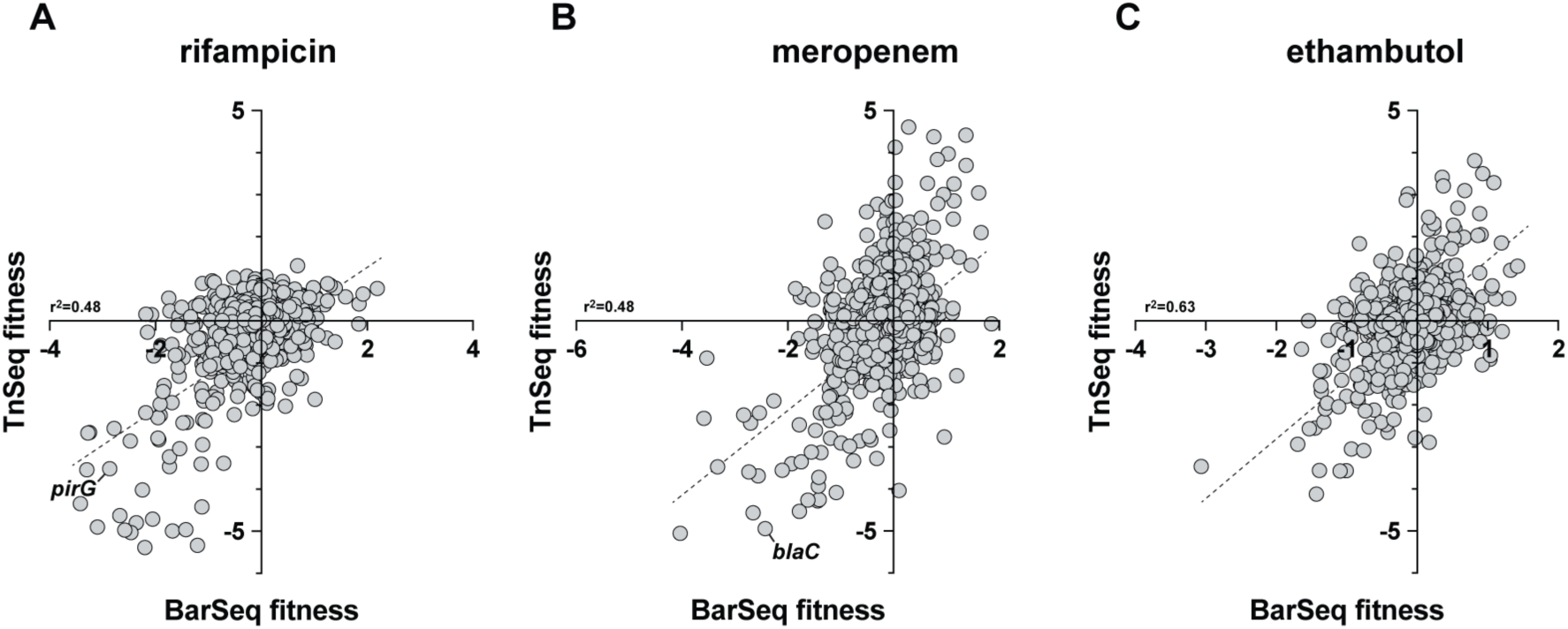
TnSeq vs RB-TnSeq. Previously published TnSeq fitness [28] plotted against RB-TnSeq fitness for rifampicin, meropenem and ethambutol. Dotted line and r^2^ represent linear correlation between statistically significant hits from RB-TnSeq with TnSeq.

**S3 Figure.**
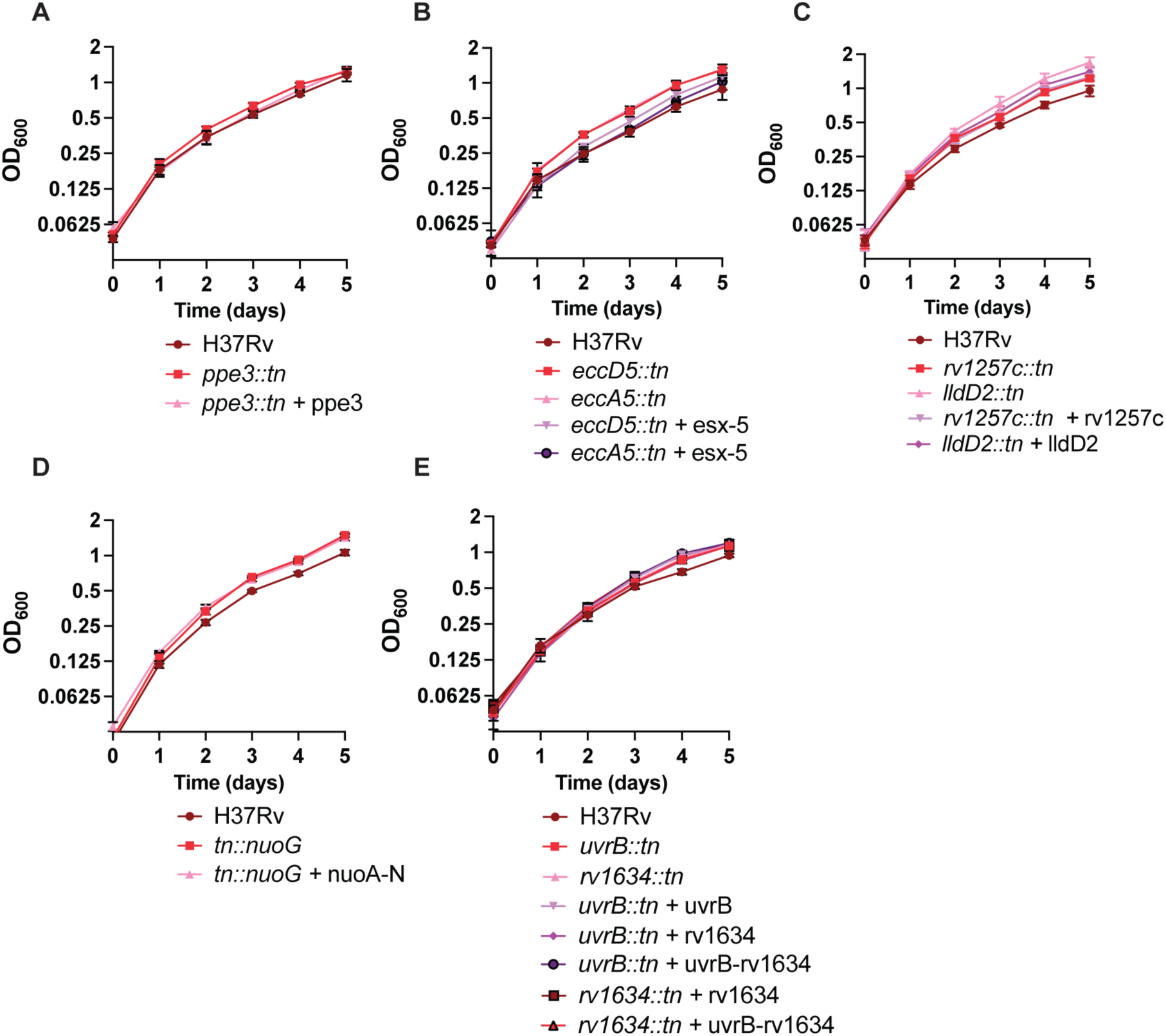
Growth curves of transposon mutants in 7H9. Growth of transposon mutants and complemented strains in 7H9 measured by OD^600^ over the course of 5 days. Data represent mean ± sd for *n*=6 replicates from two independent experiments.

**S4 Figure.**
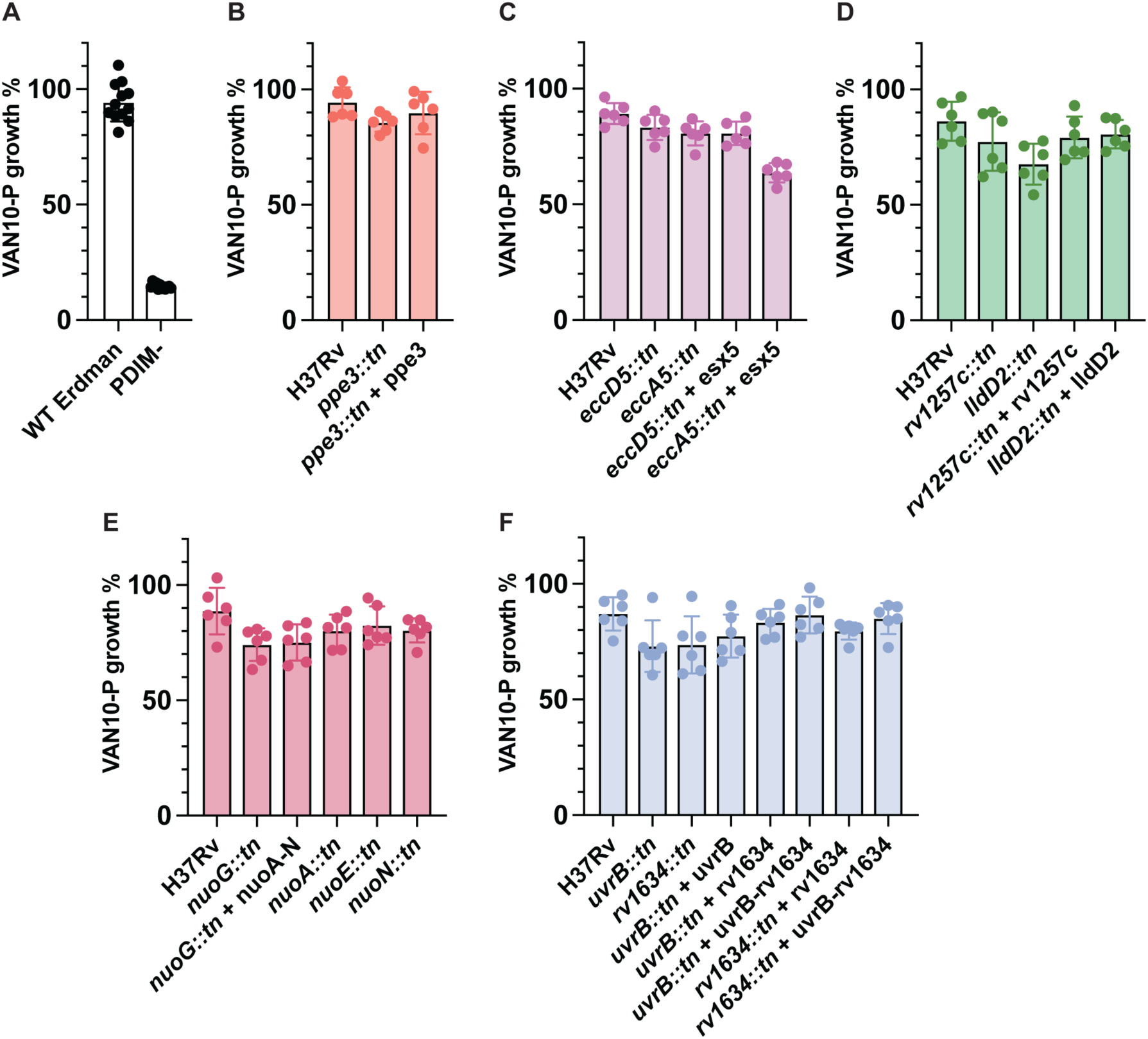
Van-10-P assay with of transposon mutants and complements. Van- 10-P assay as a proxy for the PDIM levels for strains used in this study. WT Erdman and an Erdman strain lacking PDIM production (PDIM-) are controls. Data represent mean ± sd for at least *n*=6 replicates from two independent experiments.

**S5 Figure.**
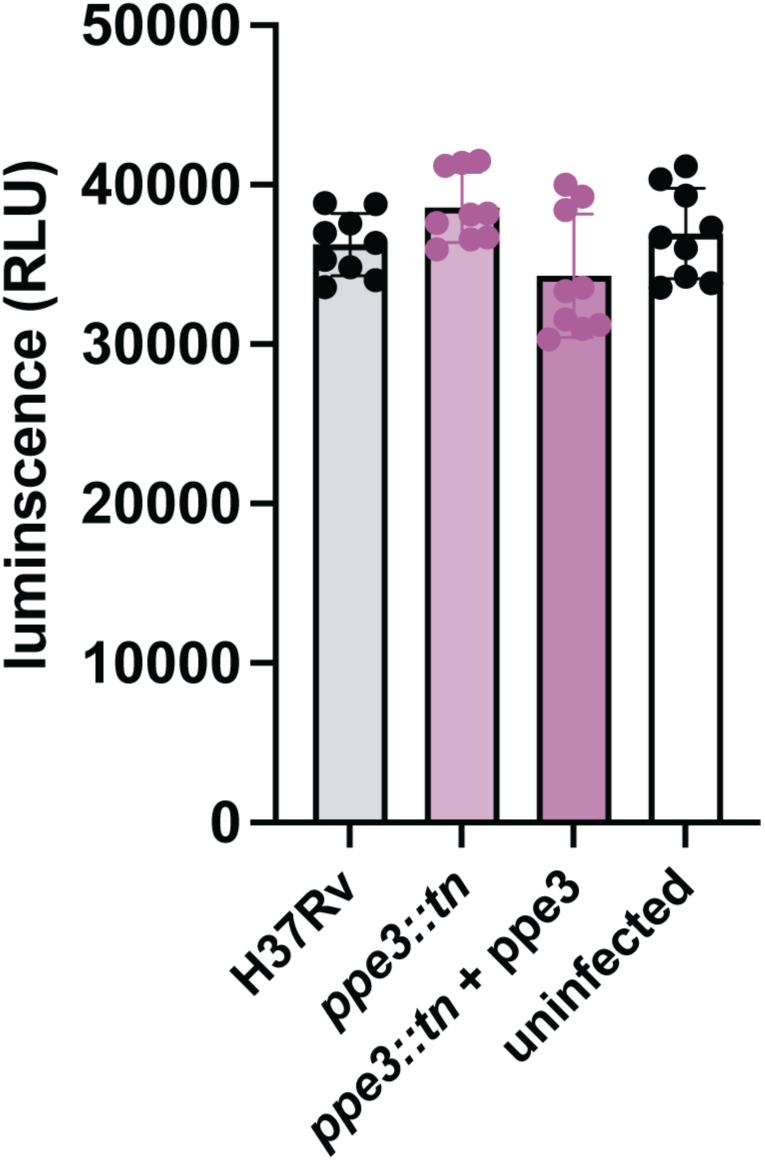
Cell viability during Mtb infection of mouse bone marrow-derived macrophages. Bone marrow-derived macrophage viability measured by cell titer glo on day 4 after infection with H37Rv, *ppe3::tn,* and ppe3 complemented strain. Data represent mean ± sd for *n*=9 replicates from three independent experiments.

**S6 Figure.**
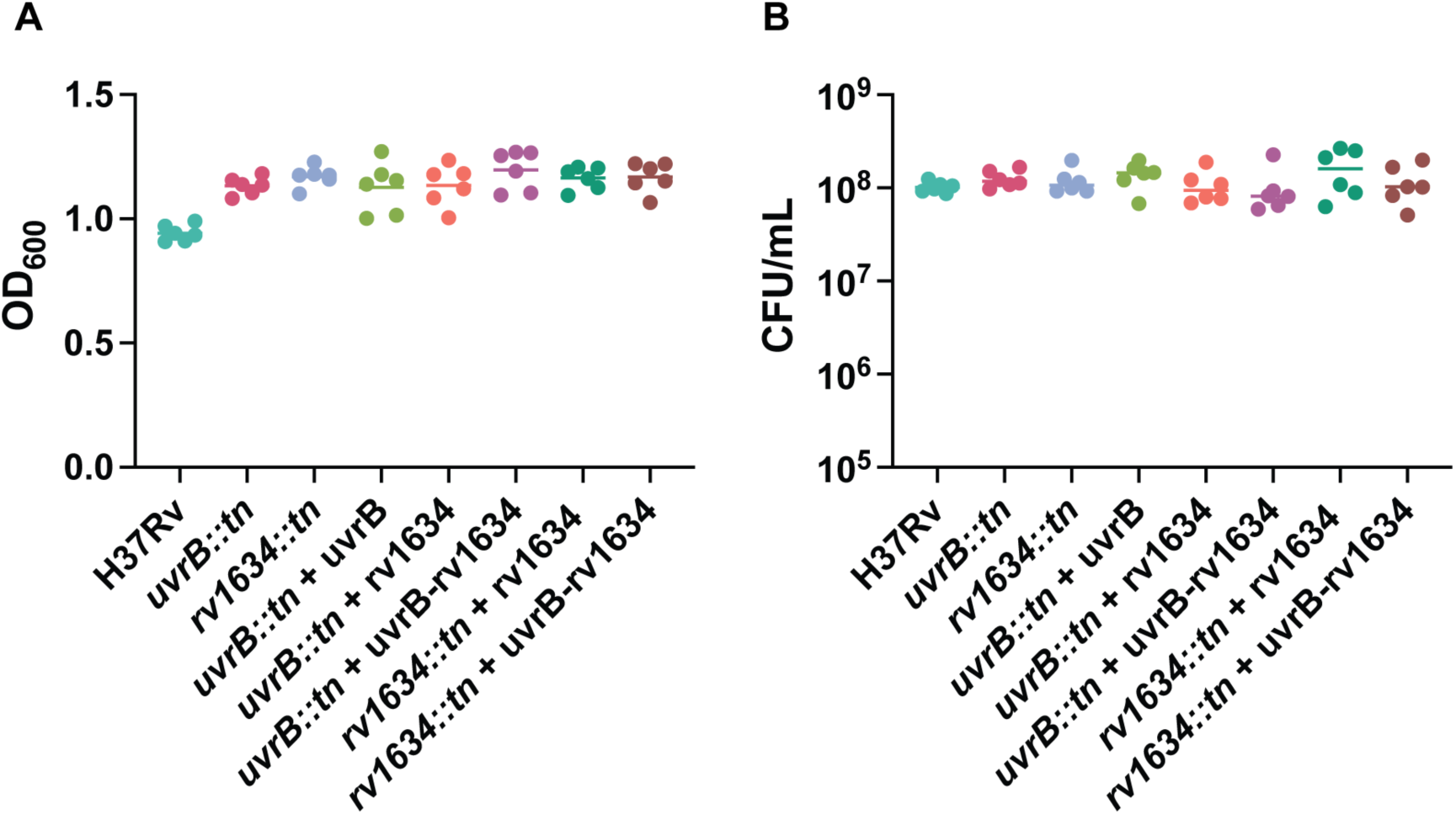
Untreated controls of pretomanid-resistant strains and complements. (A) OD^600^ and (B) CFU of untreated H37Rv, *uvrB::tn,* or *rv1634::tn* and complemented strains on day 5. Data represent mean ± sd for *n*=6 replicates from two independent experiments.

